# Deciphering the genic basis of environmental adaptations by simultaneous forward and reverse genetics in *Saccharomyces cerevisiae*

**DOI:** 10.1101/087510

**Authors:** Calum J. Maclean, Brian P.H. Metzger, Jian-Rong Yang, Wei-Chin Ho, Bryan Moyers, Jianzhi Zhang

## Abstract

The budding yeast *Saccharomyces cerevisiae* is the best studied eukaryote in molecular and cell biology, but its utility for understanding the genetic basis of natural phenotypic variation is limited by the inefficiency of association mapping owing to strong and complex population structure. To facilitate association mapping, we analyzed 190 high-quality genomes of diverse strains, including 85 newly sequenced ones, to uncover yeast’s population structure that varies substantially among genomic regions. We identified 181 yeast genes that are absent from the reference genome and demonstrated their expression and role in important functions such as drug resistance. We then simultaneously measured the growth rates of over 4500 lab strains each deficient of a nonessential gene and 81 natural strains across multiple environments using unique DNA barcode present in each strain. We combined the genome-wide reverse genetic information with genome-wide association analysis to determine potential genomic regions of importance to environmental adaptations, and for a subset experimentally validated their role by reciprocal hemizygosity tests. The resources provided permit efficient and reliable association mapping in yeast and significantly enhances its value as a model for understanding the genetic mechanisms of phenotypic polymorphism and evolution.

## INTRODUCTION

Understanding the genetic underpinnings of phenotypic variation is a major goal of modern biology. Model organisms play a prominent role in this endeavor because of the wealth of accumulated biological information and tools available for manipulating and examining these organisms. The budding yeast *Saccharomyces cerevisiae* has long been a favored eukaryotic model organism to molecular and cell biologists and was the first eukaryote to have its genome fully sequenced^1^. Large-scale phenotyping of gene deletion^2–4^ and overexpression^5,6^ strains has provided extensive data on gene function, while the availability of genome sequences from closely related species^7–10^ offers evolutionary insights into genotype-phenotype mapping. However, yeast studies have largely focused on a few laboratory strains that are phenotypic outliers^11^. To combat this issue, research on the natural diversity and ecology of *S. cerevisiae* has intensified in recent years. *S. cerevisiae* has now been isolated globally from diverse natural and man-made environments^12,13^, each of which presents potentially unique challenges for growth and survival. Therefore, *S. cerevisiae* offers a rich system in which to investigate the genetic basis of environmental adaptation. For comparison, although *S. cerevisiae*’s genomic diversity is ~20% that of its sister species *S. paradoxus*, its phenotypic diversity is much greater^11^.

Studying the genetic basis of yeast environmental adaptation requires knowledge about its population structure and genomic diversity. Such knowledge has accumulated primarily through low-coverage Sanger sequencing^12^ and tiling array hybridization^14^. These studies, as well as restriction-site associated DNA sequencing of a large strain set^15^, have revealed a complex population structure of the species. High-quality genomes produced by next-generation sequencing further revealed copy-number variations and genomic rearrangements in the species^16–18^ as well as the origins of domestic *S. cerevisiae* strains^19,20^. However, because of human activity, many *S. cerevisiae* strains have complex ancestry from multiple lineages and are mosaics while strains representing pure lineages are often phenotypically distinct^11^. This strong and complex population structure has made genome-wide association study (GWAS), an important forward genetic method for detecting influential genetic variants in many species, difficult in yeast^21,22^, hindering the use of an otherwise powerful model species for systematic analysis of the genetic basis of phenotype variations. To overcome this hurdle, we have developed a resource for efficient GWAS in *S. cerevisiae*. Specifically, we generated high-quality genome sequences of 85 diverse *S. cerevisiae* strains that represent genetic and phenotypic variations of the species. Combining them with the available high-quality genome sequences in the literature, we assembled a dataset of 190 genomes. We then conducted a comprehensive population genomic analysis, including elucidating detailed phylogenetic relationships of the strains and population structure of the species. We detected genes from the newly sequenced genomes that are absent from the reference genome and demonstrated their expression and functions. We barcoded diverse natural strains, allowing simultaneous phenotyping of them with over 4500 single gene deletion strains by a high-throughput barcode-sequencing (bar-seq) method^23^. With the assistance of reverse genetics, our GWAS identified potential causal genes responsible for growth rate variations in five of six environments examined. We experimentally verified a subset of these associations for high-temperature growth by a reciprocal hemizygosity test^24^, establishing this approach as a powerful method for unbiased identification of the genetic basis underlying environmental adaptation in *S. cerevisiae*.

## RESULTS

### Population genomics of *S. cerevisiae*

To identify genetic variation underlying phenotypic variation among *S. cerevisiae* strains, we generated high-quality genome sequences of 85 strains collected in six continents and from a variety of human associated and wild environments (**Fig. 1a, Supplementary Data 1**). We obtained an average of 3.75 million 2×100-nucletide paired-end reads per strain, approximately 97% of which were successfully mapped to the S288c reference genome. This resulted in an average coverage of 60× per genome (range 38-99×) (**Supplementary Data 2**). On average, 6% of the reference genome was not covered by a read in each sequenced strain due to stochastic sampling of reads and/or strain differences in gene content as well as repeat elements. In total, we identified 311,287 single nucleotide polymorphisms (SNPs) and 15,884 insertions/deletions (indels).

**Fig. 1.**
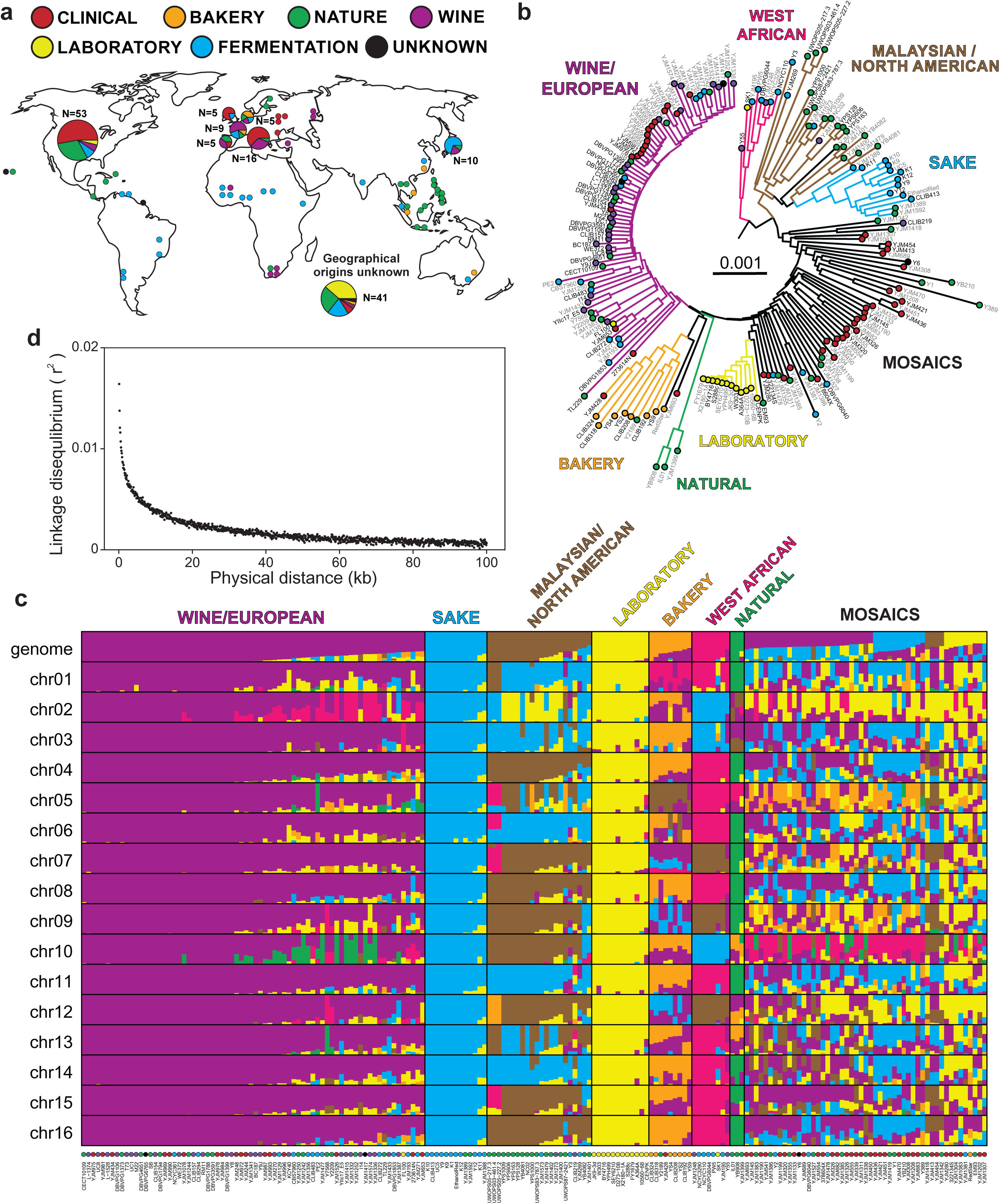
Geographical, environmental, and phylogenetic relationships of the 190 *S. cerevisiae* strains analyzed. (**a**) World map indicating the geographic locations where the analyzed strains were sampled. Colors represent the environment of isolation if known. (**b**) Maximum composite likelihood neighbor-joining tree of the 190 strains based on genome-wide SNP data. The environment type from which each strain was isolated is indicated if known along with the clades into which the strains have been assigned. The scale bar represents 0.1% genome sequence divergence. Strain names in black are those sequenced in this work while grey were previously sequenced. The same tree with bootstrap values is shown in Supplementary Fig. 1. Trees based on individual chromosomes are provided in Supplementary Fig. 2. (**c**) Population structures of the 190 strains inferred based on the entire genome or individual chromosomes. Strains are arrayed based on clade membership in panel (b), with colored circles indicating the environmental origins of the strains. Different colors show different inferred populations, which are indicated at the top of the panel. The Y-axis shows the fraction of SNPs coming from each inferred population. (**d**) Linkage disequilibrium (LD) decays as the physical distance between two linked sites becomes larger. LD is measured by *r*^2^ between two linked sites minus the mean *r*^2^ between two sites located on different chromosomes.

To acquire a comprehensive view of *S. cerevisiae* strain relations, we identified 105 additional strains that have publically available high-quality genome sequences^16,17^ at the time of our analysis (August 2016), and applied our analysis pipeline to the Illumina sequencing reads of these strains. A neighbor-joining tree of all 190 strains was then constructed on the basis of a combined set of 421,773 SNPs (**Fig. 1b, Supplementary Fig. 1, Supplementary Data 1, 2**). The tree was rooted using outgroup sequences from recently identified Chinese isolates^13^. We recovered phylogenetic clustering based roughly on geographical and environmental origins of the strains, consistent with early observations made from fewer strains and SNPs^12,14^. Clustering can be seen of strains into the West African, Malaysian/North American, Sake, Laboratory, and European/Wine groups previously identified. Additionally, we identified a “Bakery” clade that was previously unrecognized and a new clade of three wild strains (one from soil in Illinois and two on the gums of wild cherry trees from unknown locations) (**Fig. 1b**). The remaining strains, originating from a wide variety of environments, form a paraphyletic group named “Mosaics” (**Fig. 1b**; see below).

To examine the population structure of *S. cerevisiae*, we employed a model-based clustering algorithm implemented in fastSTRUCTURE^25^, and identified seven distinct subpopulations (top row in **Fig. 1c**) that are in agreement with the strain isolation sources and corroborate the clustering pattern seen in the phylogeny. In addition, many strains from the paraphyletic group mentioned are mosaics with ancestry from several lineages (top row in **Fig. 1c**), supporting previous observations based on smaller data^12^. We then conducted the fastSTRUCTURE analysis for each of the 16 chromosomes and observed prominent among-chromosome variations in population structure (**Fig. 1c**). For example, the West African subpopulation is genetically distinct from other subpopulations for 10 of its 16 chromosomes, but is indistinct from the sake subpopulation for three chromosomes and indistinct from the Malaysian/North American subpopulation for another three chromosomes. This variation in chromosomal population structure is indicative of differences in the evolutionary histories of different chromosomes due to pervasive gene flow. Because *S. cerevisiae* reproduces largely asexually^26^, rare crosses between lineages can establish unique populations where distinct chromosome combinations persist in the absence of outbreeding. The observed differences in chromosomal population structure are not due to stochasticity in the population structure assessment because multiple runs on the same chromosome displayed only minor variations. Furthermore, phylogenies reconstructed using SNPs from individual chromosomes corroborate the fastSTRUCTURE results (**Supplementary Fig. 2**).

The extent of linkage disequilibrium (LD) between SNPs is an important characteristic determining the highest possible resolution of association analysis; the lower the LD, the higher the resolution can be. We found that the mean LD measured by *r*^2^ equals 0.0164 for SNPs within 100 nucleotides and it halves as the physical distance increases to 1200 nucleotides (**Fig. 1d**). Given that the average distance between the beginning of one gene and that of the next gene on the chromosome is ~2kb in yeast, this result indicates that fine-scale mapping to the gene level is possible by GWAS in this species.

To further characterize the genetic variation present in *S. cerevisiae*, we conducted a comprehensive population genomic analysis of all 190 strains. The basic population genetic parameters are summarized in **Table 1**. Intronic and intergenic polymorphisms (θ_W_) and nucleotide diversities (π) are significantly lower than those at synonymous sites, suggesting pervasive purifying selection acting on noncoding regions. This is consistent with the fact that the compact nature of the yeast genome results in intergenic regions dense with promoter and other important regulatory elements and that yeast introns can regulate gene expression^27,28^. Consistent with previous reports using smaller data^12^, no positive selection was detected by virtually every population genetic test applied, and the estimated fraction of adaptive amino acid substitutions was 0 (**Supplementary Figs. 3-5, Supplementary Tables 1, 2**; see Methods).

**Table 1.** Polymorphism (*θ*_W_) and nucleotide diversity (*π*) per kb in different regions of the yeast genome

| | $\theta_w$ | | $\pi$ | |
| --- | --- | --- | --- | --- |
|  | Mean | SD | Mean | SD |
| Intergenic | 5.63 | 0.018 | 2.67 | 0.015 |
| Intronic | 7.52 | 0.135 | 3.68 | 0.122 |
| Synonymous | 14.60 | 0.035 | 9.08 | 0.037 |
| Nonsynonymous | 3.63 | 0.010 | 1.37 | 0.007 |
| Total | 5.97 | 0.009 | 2.99 | 0.009 |

Finally, for the 85 newly sequenced genomes, we also analyzed and observed intron presence/absence polymorphisms (**Supplementary Fig. 6**; see Methods) and identified aneuploidies and large segmental duplications (**Supplementary Fig. 7**; see Methods).

### Non-reference genes

Our knowledge of *S. cerevisiae* genome content and function is largely derived from a few laboratory strains, which are now known to be phenotypically atypical^11^. Furthermore, as S288c, the strain from which the reference set of genes are defined, has long been maintained in relatively benign and unvarying laboratory environments, the possibility arises that genes important for survival only in non-laboratory conditions may have been lost. Additionally, the isolation of yeast strains from disparate environments suggests that the local ecology with which they interact is also unique and that horizontal gene transfer (HGT) from other species within their local environment may introduce new genes. Due to their potential importance in understanding a strain’s phenotype, identifying coding regions not present in the reference strain is of great interest. To this end, we performed a *de novo* genome assembly of the Illumina sequencing reads obtained from our 85 strains (see **Supplementary Data 3** for assembly statistics). We identified 181 distinct non-reference genes distributed across the phylogeny (**Fig. 2a, Supplementary Fig. 8, Supplementary Data 4**). The majority were found to have a BLAST hit in a previously sequenced non-S288c *S. cerevisiae* strain, while others had best hits in other fungi or more distantly related organisms (**Fig. 2a**). The phylogenetic distribution of non-reference genes did not follow a straightforward pattern; distantly related strains often share non-reference genes, suggesting independent gains via HGT or introgression, or independent losses (**Fig. 2a, Supplementary Fig. 8).**

**Fig. 2.**
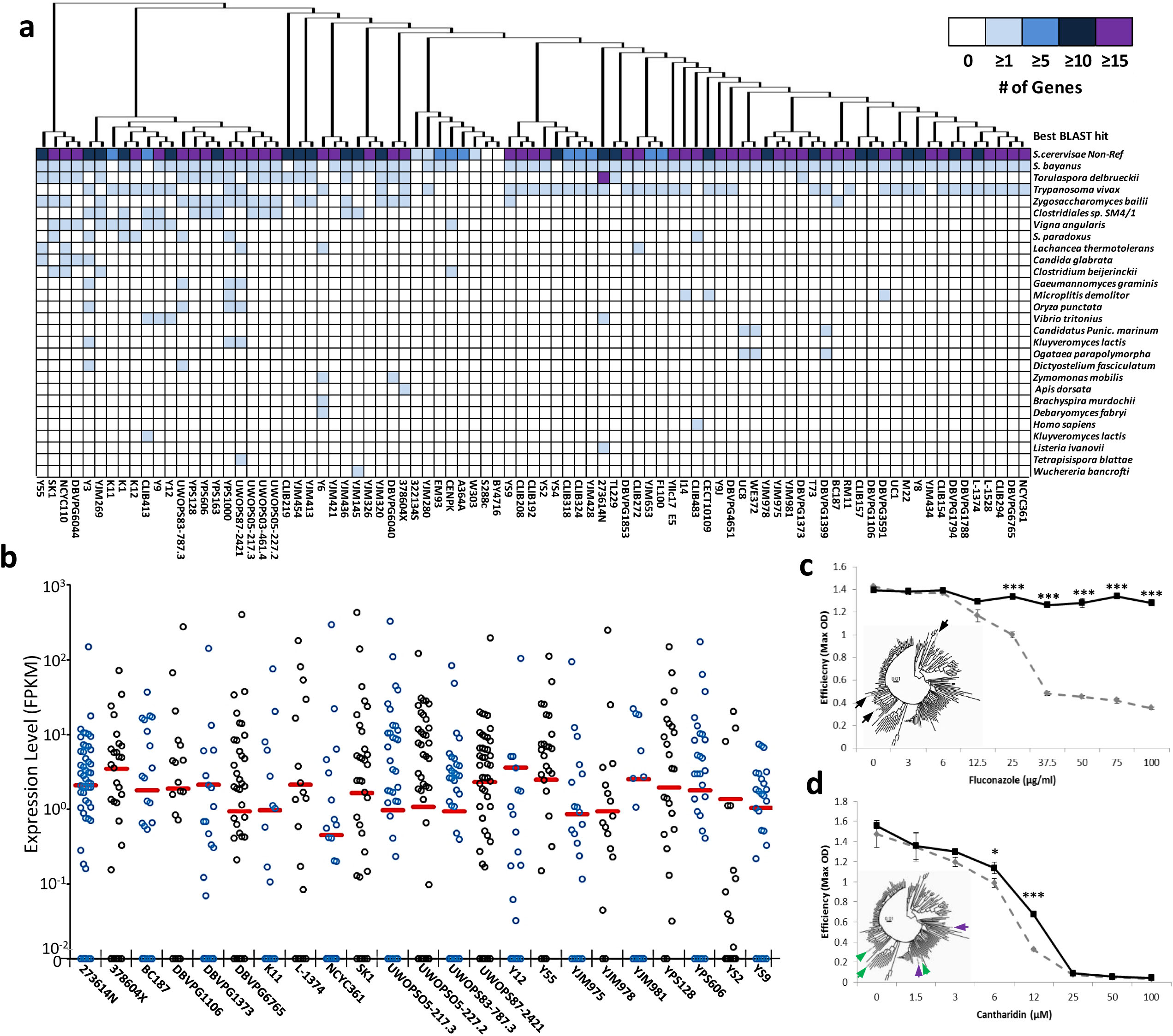
Origins, expressions, and functions of non-reference genes identified from the 85 newly sequenced *S. cerevisiae* genomes. (**a**) Evolutionary origins of non-reference genes. Color indicates the number of non-reference genes identified from each of the 85 genomes (strain name shown at the bottom of the panel) that received the best hit in a particular species listed at the right-hand side of the panel. The relationships of the 85 strains, as in Fig. 1b, are indicated by the phylogeny. (**b**) Expression levels of non-reference genes estimated using existing mRNA sequencing data for 23 of the 85 sequenced strains. Each circle represents a non-reference gene in a strain indicated at the bottom of the panel. The red horizontal bar represents the lower fifth percentile of gene expression levels of all reference genes in that strain. FPKM, Fragments Per Kilobase of transcript per Million mapped reads. (**c**) Fitness consequence of deleting *Non-Ref-1* from UWOPS87-2421 in the presence of the antifungal fluconazole. Fitness is quantified by efficiency (maximum OD). Wild-type and deletion strain data are shown by black solid line and dashed grey line, respectively. *P*-values from *t*-tests of the null hypothesis of no fitness effect from the gene deletion are indicated as follows: *, ≤ 0.05; **, ≤ 0.01; ***, ≤ 0.001. Error bars represent the standard error of the mean from three replicates. The inset shows the same tree as in Fig. 1b, with the black arrows pointing to the strains where *Non-Ref-1* is found. (**d**) Fitness consequence of deleting *Non-Ref-2* from CLIB272 in the presence of the antifungal cantharidin. All notations are the same as in panel (c). The inset shows the same tree as in Fig. 1b, with the green arrows pointing to the three strains carrying the first variant (CLIB272, YJM1326 and YJM428) and the purple arrows pointing to the two strains carrying the second variant (Y6 and YJM1355).

We estimated the expression levels of these non-reference genes present in 23 *S. cerevisiae* stains with available transcriptome data generated by mRNA sequencing (RNA-seq)^29^. We found that on average, 54.9% of the non-reference genes examined had a higher expression level than each of the 5% most lowly expressed reference genes (**Fig. 2b, Supplementary Data 5**). Because the RNA-seq data were collected in a single benign environment, it is likely that, due to conditional expression, more of the non-reference genes identified are expressed at appreciable levels in the appropriate environments.

To examine functional importance of non-reference genes, we focused on two of them for which our initial BLAST search revealed their closest hit to be within the well annotated *Lanchea thermotolerans* genome, suggesting HGT as opposed to introgression from a closely related *sensu stricto* species with which *S. cerevisiae* is able to mate. We used growth curves to determine the phenotypic consequences of deleting these non-reference genes on two aspects of strain growth, maximum growth rate and efficiency (see Methods). *Non-Ref-1*, identified from the Malaysian strain UWOPS87-2421, resembles the *L. thermotolerans* coding region KLTH0E00528g, which is in turn annotated as a homolog of *S. cerevisiae FLR1*, a multi-drug transporter responsible for the efflux of drugs such as the widely used anti-fungal fluconazole^30^. Thus, *Non-Ref-1*, expressed even in a benign environment (5.25 RPKM; **Supplementary Data 5**), may confer resistance to this important drug. We deleted *Non-Ref-1* from haploid UWOPS87-2421 cells and exposed both the wild-type and deletion strains to various fluconazole concentrations to investigate the impact of gene deletion on strain growth (**Fig. 2c**). Deleting *Non-Ref-1* has a small but significant effect on the maximum growth rate (**Supplementary Fig. 9a**) and a large effect on growth efficiency especially when fluconazole concentration exceeds 25µg/ml, which is typical in high-dose fluconazole clinical treatment^31,32^ (**Fig. 2c**). If *Non-Ref-1* is indeed a drug transporter similar to *FLR1*, it may also be involved in resistance to diazaborine, benomyl, methotrexate, and other drugs^33,34^.

To better understand the history of *Non-Ref-1*, we performed additional BLAST searches in other published *S. cerevisiae* genomes that were built *de novo*^17^. We found in YJM653 an intact *Non-Ref-1* and in YJM1250 an apparently pseudogenized *Non-Ref-1* that is disrupted by an insertion; these two strains respectively reside at the edge of and within the Wine/European clade, both being highly diverged from UWOPS87-2421 (**Fig. 2c**). UWOPS87-2421 and YJM653 differ at only one nonsynonymous site and no synonymous site in this 1644-nucleotide gene. The highly similar chromosomal locations of all three *Non-Ref-1* genes in an unstable telomeric region (UWOPS87-2421, ChX:33185-34826; YJM653, ChX:30819-32462; YJM1250, ChX:22819-24457) suggest a single origin of *Non-Ref-1* in *S. cerevisiae*.

The second non-reference gene experimentally studied, *Non-Ref-2*, was initially identified in strains CLIB272 and Y6 and found to be similar to the *L. thermotolerans* gene *KLTH0H09460g*, which is homologous to *CRG1* in *S. cerevisiae*, a methyltransferase gene involved in lipid homeostasis and providing resistance to the phosphatase inhibitor cantharidin^35^. Unlike *Non-Ref-1, Non-Ref-2* is not expressed in benign conditions (**Supplementary Data 5**). However, because *CRG1* expression increases by 40-50 fold upon exposure to cantharadin and other stresses^35^, it is possible that *Non-Ref-2* expression is condition-specific. We deleted *Non-Ref-2* from haploid CLIB272 cells and exposed wild-type and deletion strains to varying concentrations of cantharadin (**Fig. 2d**; see Methods). The deletion of *Non-Ref-2* does not alter the maximum growth rate consistently (**Supplementary Fig. 9b**), but has a significant effect on growth efficiency at intermediate drug levels, with the deletion strain reaching only ~50% of the maximum OD of the wild-type strain at a cantharadin concentration of 12µM (**Fig. 2d**). Additional BLAST searches detected *Non-Ref-2* in five *S. cerevisiae* genomes previously published^17^. We identified two distinct versions of *Non-Ref-2* distributed across the phylogeny (inset of **Fig. 2d**) that differ at 10 sites, including four nonsynonymous sites. However, the similar genomic location of these two variants, in a telomeric region of Ch. XV, suggests a single acquisition followed by divergence and outcrossing to result in their current phylogenetic distribution.

### High-throughput simultaneous strain phenotyping by bar-seq

Accurate phenotyping is central to uncovering the genetic basis of phenotypic variation. Phenotyping different natural yeast strains has primarily relied on the production and analysis of growth curves^11^ or digital photography-based colony sizes^36^, which have limited throughput and resolution. We decided to adopt bar-seq^23^ for phenotyping, which measures the growth rates of all strains of interest in the same test tube by Illumina sequencing of strain-specific DNA barcodes, substantially increasing the throughput and resolution of growth rate estimation. Bar-seq was originally designed to quantify the relative growth rates of S288c-derived gene deletion strains each carrying two unique pieces of 20-nucleotide DNA (barcodes) inserted at the time of strain construction^4,37^. We similarly constructed a panel of natural strains, each of which carries two unique barcodes which are not present in any deletion strain. This not only allows the bar-seq experiment of the natural strains but also that of natural strains and deletion strains all in one test tube (**Fig. 3a**).

**Fig. 3.**
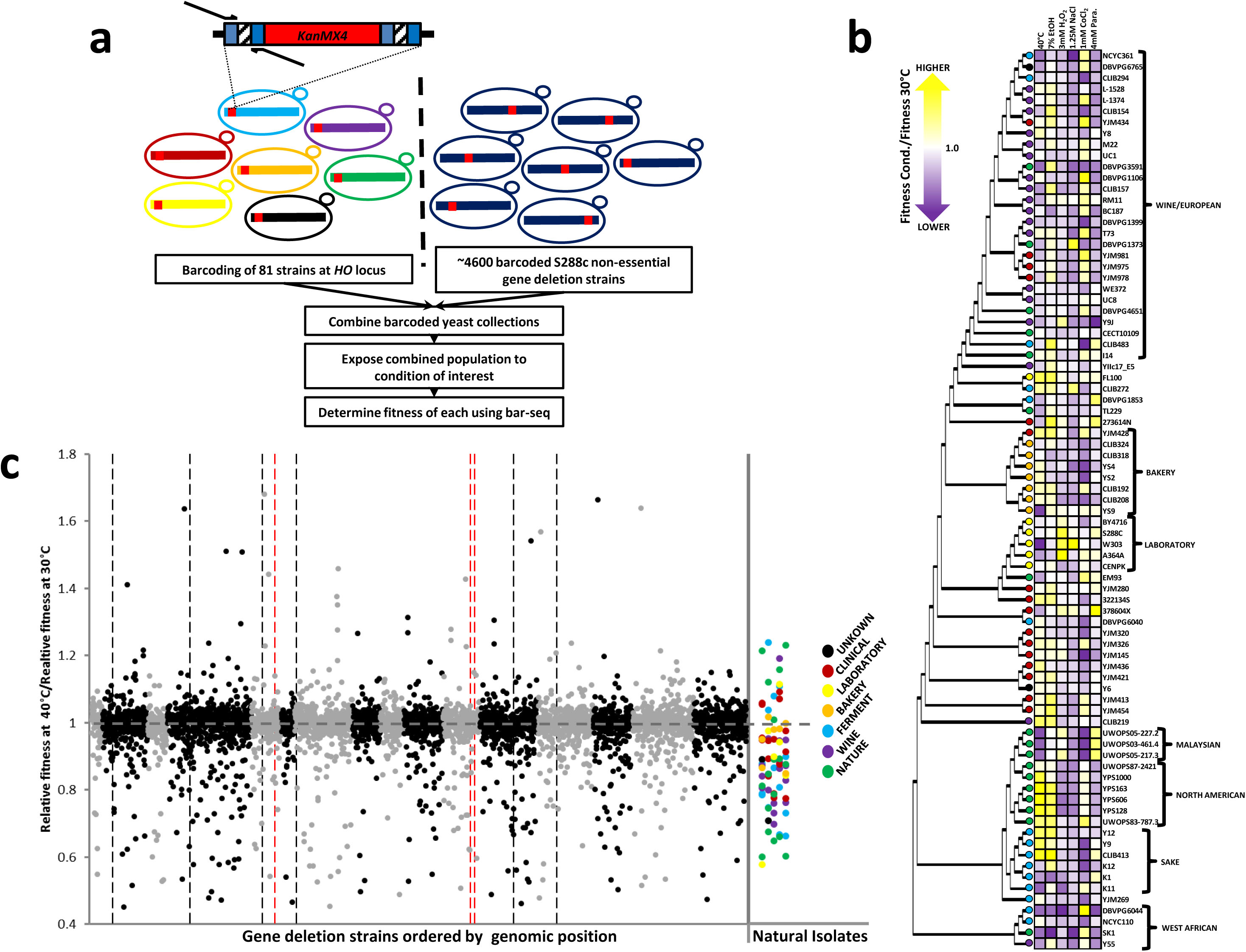
Simultaneous high-throughput phenotyping of 81 natural isolates and 4521 gene deletion strains. (**a**) Flow chart showing the procedure of barcoding the natural isolates and simultaneously phenotyping natural isolates and gene deletion strains. (**b**) Heat map showing the fitness of 81 natural isolates relative to the reference strain (S288c-derived *HO* deletion strain) in each stressful environment, relative to that in the benign environment of YPD at 30°C. The colored arrows show the color scheme for relative fitness, with the most extreme colors depicting the most extreme fitness values in each environment. Colored circles indicate the environmental origins of the natural isolates as in Fig. 1a. (**c**) The fitness of 81 natural isolates and 4521 S288c derived gene deletion strains relative to the reference strain at a high temperature environment, relative to that in the benign environment. The natural isolates are shown by colored circles, based on the color scheme in Fig. 1a. The gene deletion strains are shown by black or grey circles depending on their locations on odd-numbered or even-numbered chromosomes, respectively, and are arranged by chromosomal position. Dashed vertical lines indicate the regions identified as significant by GWAS with those shown in red denoting those further investigated by hemizygozity tests in Fig. 4.

We successfully inserted the unique barcodes, flanking a G418 sulfate resistance marker (*KanMX4*), at the *HO* (YDL227C) locus of 81 of the 85 diploid strains sequenced here (**Supplementary Fig. 10, Supplementary Data 1**; see Methods). Of these 81 heterozygous *HO/ho::KanMX4* strains, we obtained stable a and α haploids for 76 of them before also obtaining a *ho::HygMX4* (Hygromycin B) and α *ho::NatMX4* (nourseothricin resistance) haploids through marker switching (**Supplementary Data 1**; see Methods).

To test the utility of these barcoded strains for studying the genetic basis of phenotypic variation, we combined the 81 barcoded diploid strains with the *S. cerevisiae* nonessential gene deletion collection constructed from the S288c background to create a common starter pool to use across our experiments (**Fig. 3a**). We then subjected this pool to a benign environment (YPD at 30°C) as well as six stressful environments: high temperature (YPD at 40°C), high salt/osmotic stress (YPD at 30°C + 1.25M NaCl), high ethanol (YPD at 30°C + 7% ethanol), superoxide anions (YPD at 30°C + 4 mM paraquat), oxidizing agents (YPD at 30°C + 3 mM hydrogen peroxide), and a hypoxia mimetic (YPD at 30°C + 1mM cobalt chloride). We extracted genomic DNAs from the common starting pool and following each competition, produced bar-seq libraries, and quantified the sequencing read number of each barcode, a proxy for strain frequency, at each time point, by Illumina sequencing (**Fig. 3a**). We estimated the growth rate or fitness of each strain in a particular environment relative to that in the benign 30°C YPD environment, relative to the corresponding value of a reference strain (S288c-derived *HO* deletion strain) to identify strains with particularly high or low relative fitness in the environment of interest. **Fig. 3b** shows these relative fitness values for the 81 strains for each of the six stressful environments. While there is a clear phylogenetic component to the phenotypic similarity among some strains, being particularly apparent within the North American and Malaysian clades, other regions appear phenotypically diverged from the strains they are most closely related to (**Fig. 3b**). The bar-seq data also allowed us to determine the effect each gene deletion has across the tested environments. **Fig. 3c** plots the fitness under a high temperature challenge relative to the benign environment compared to the reference strain for over 4500 BY strains each deficient of a nonessential gene as well as the 81 natural strains. The relative fitness varies greatly among the gene deletion strains, with 13.8% (623/4521) of gene deletion strains having significantly higher relative fitness and 18.3% (829/4521) having significantly lower relative fitness than the reference strain (**Fig. 3c, Supplementary Data 6**). Generally similar patterns are also observed in the other five environments examined (**Supplementary Fig. 11, Supplementary Data 6**).

### Identifying phenotypically relevant SNPs by GWAS

Using a multi-stage GWAS approach (**Supplementary Fig. 12**; see Methods), we attempted to identify SNPs associated with relative fitness variations among the 81 natural strains. We discovered between 3 and 19 associated SNPs per stressful environment after controlling for population structure, for five of the six environments examined (**Supplementary Data 7**). Due to LD, an associated SNP may not be a causal genetic variant. Although we have no direct evidence that the detected SNPs, or indeed linked regions, are responsible for the phenotypic variations observed, a closer inspection of the regions surrounding the associated SNPs suggests that most signals are real. For example, we detected 13 SNPs associated with relative fitness at 40°C. Five of the 13 SNPs map to a ~16 kb region on Ch. XI that contains the ribosomal protein gene *RPS21a*, deletion of which is known to slow growth at high temperature (www.yeastgenome.org). Similarly, a second SNP on Ch. XI, located 51.5 kb from this cluster, is within 7 kb upstream and downstream of the genes *DBP7, RPC37* and *GCN3*, which again are known to reduce heat tolerance upon deletion, as well as *SET3* and *YKR023C*, known to reduce stress tolerance when deleted.

While association studies rarely validate that identified SNPs or linked regions are responsible for the observed phenotypic differences, such validation can be performed in yeast. To this end, we used reciprocal hemizygozity tests to identify difference in fitness due to deletion of alternate alleles in hybrids of high- and low-fitness strains at 40°C relative to 30°C for genes surrounding each of several associated SNPs (**Fig. 4a**; see Methods). In each competition experiment, one strain expressed yellow florescent protein (YFP), facilitating the quantification of relative fitness by flow cytometry^38^. Reciprocal experiments with the YFP marker in opposing hybrid background were performed to remove any fitness effect of YFP expression. We chose two strains with high fitness (YPS128 and YJM320) and two with low fitness (W303, UWOPS05-227.2) at 40°C relative to 30°C and selected three significant SNPs for investigation, reciprocally deleting four to six genes surrounding each SNP (**Supplementary Data 7**).

**Fig. 4.**
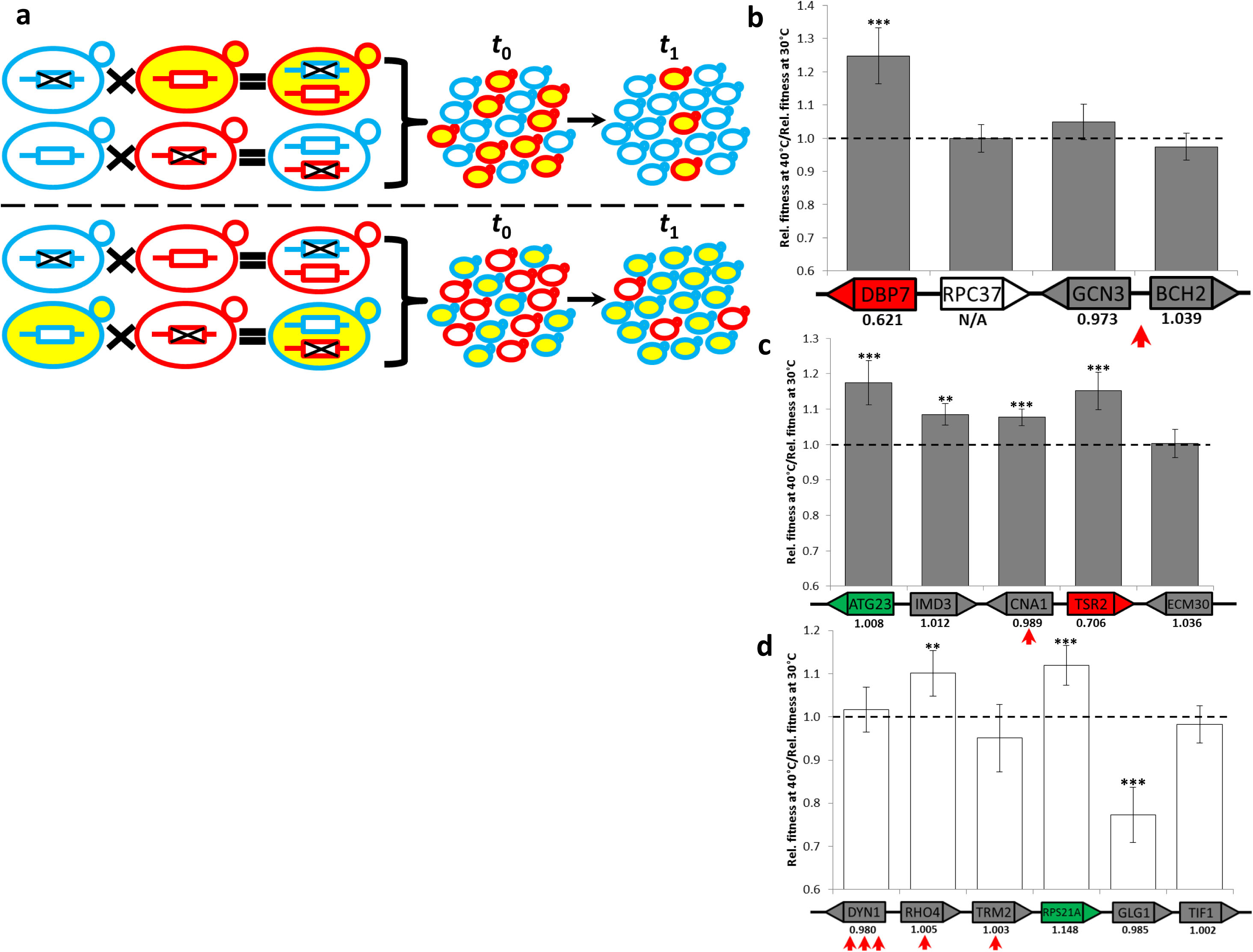
Reciprocal hemizygosity test of causal effects of candidate genes surrounding SNPs identified by GWAS to be associated with specific adaptations to a 40°C environment. (**a**) Flow chart depicting the procedure of the reciprocal hemizygosity test. The blue- and red-outlined cells respectively depict the initial haploid cells carrying the predicted high- and low-fitness alleles, as well as the hemizygous cells carrying the predicted high- and low-fitness alleles. Each black cross indicates a gene deletion, and yellow colored cells indicate YFP expression. The competitions above and below the horizontal line have alternative YFP-marked genotypes, allowing removal of the potential fitness effect of YFP expression on our assay. (**b**) Fitness of the hemizygous strain deficient of the low-fitness allele of a candidate gene relative to that deficient of the high-fitness allele in the high-temperature environment, relative to that in the benign environment. The low- and high-fitness alleles are from W303 and YPS128, respectively. Significant deviation of fitness from 1 is determined by a *t*-test using biological replicates and is indicated as follows: *, *P* ≤ 0.05; **, *P* ≤ 0.01; ***, *P* ≤ 0.001. Error bars denote the 95% confidence intervals determined by Fieller’s theorem. The genes examined are shown at the bottom of the panel, with the red arrow pointing to the associated SNP detected from GWAS. The number below each gene is the fitness of the gene deletion strain relative to that of the reference strain at 40°C relative to 30°C, as shown in Fig. 3c. Genes with significant positive effects on relative fitness (when deleted) are indicated in green, while significant negative effects are indicated in red. Additionally, grey and white coloring of genes denote no significant effect and no data available, respectively. (**c**) Same as panel (b) except for a different genomic region. (**d**) Same as panel (b) except for a different genomic region and the strains used. The low- and high-fitness alleles are from UWOPS05-227.2 and YJM320, respectively. Grey and white vertical bars denote different genetic backgrounds in panels (b)-(c) and panel (d).

The first SNP of interest (**Fig. 4b**), the most significant identified, is located at site 490,822 of chromosome 11 within the bidirectional promoter of *GCN3* and *BCH2*, both annotated as having temperature related deletion phenotypes. In addition, two neighboring genes (*DBP7* and *RPC37*) are similarly annotated. In particular, our bar-seq data showed that deleting *DBP7* from S288c drastically reduces the fitness at 40°C compared to that at 30°C. We respectively deleted the alternate alleles in W303/YPS128 hybrids for each of the above four genes and compared their growth rates at 40°C relative to that at 30°C. As expected, the strain retaining the predicted high-fitness allele of *DBP7* from YPS128 grew significantly faster than the strain retaining the predicted low-fitness allele from W303 at 40°C relative to 30°C (**Fig. 4b**). No such significant difference was observed for the other three genes tested (**Fig. 4b**). Thus, variation in either function or expression of *DBP7*, which encodes a putative ATP-dependent RNA helicase of the DEAD-box family, likely contributes to fitness variation at 40°C among these strains. Interestingly, the causal gene is two genes (3.5kb) away from the significantly associated SNP identified in GWAS.

The second SNP of interest is located within the coding region of *CNA1* (Ch12_1004315), a gene whose deletion is annotated as increasing stress susceptibility. The alternate alleles of *CNA1* and four neighboring genes were respectively deleted in W303/YPS128 hybrids. Unexpectedly, retention of the high-fitness allele resulted in a higher fitness than retention of the low-fitness allele at 40°C relative to 30°C for four of the five genes examined (**Fig. 4c**), suggesting that the associated SNP may reflect signals from several genes. Unlike the results for *DBP7*, deletion from S288c for two of these four genes did not have an appreciable impact on relative fitness (**Fig. 4c**). These results suggest that the genes responsible for natural phenotypic variation differ from genes affecting growth in a single laboratory strain, highlighting the necessity of considering the genetic background and type of mutation when determining the gene-phenotype relationships.

The third region investigated contained a cluster of five significantly associated SNPs on Ch. XI. Three of them are located in the coding region of *DYN1* and one in each of the coding regions of the upstream genes *RHO4* and *TRM2*. We separately deleted alternate alleles of these three genes as well as three further upstream genes including the thermally annotated gene *RPS21A* from the hybrid of UWOPS05-227.2 and YJM320. The results confirmed the causal effects of *RHO4* and *RPS21A* (**Fig. 4d**). However, deleting *RHO4* from S288c did not significantly alter the relative fitness at 40°C (**Fig. 4d**), again suggesting differences in genetic background and/or type of mutation underlying phenotypic variation in natural strains relative to a laboratory strain. Finally, for *GLG1*, the strain carrying the predicted high-fitness allele was outcompeted by the strain carrying the predicted low-fitness allele, suggesting a complex genetic architecture for growth in this environment relative to a more benign environment, not only that an associated SNP may have multiple causal genetic variants but also that these variants may have opposite fitness effects.

## DISCUSSION

We have presented here a comprehensive overview of the genomic diversity within *S. cerevisiae* by combining newly sequenced genomes with those previously published. Using this set of genomes, we thoroughly investigated the evolutionary past of these strains. The genome sequences, in conjunction with the genetically tractable haploids and diploids created, provide valuable resources to the community for understanding the genetic basis of phenotypic variation in yeast. This will not only be scientifically informative due to the wealth of biological information we have about yeast but will also be useful to society due to the wide use of diverse yeast in many industries.

We found strong population structure in *S. cerevisiae* and significant variation in population structure and evolutionary history among different parts of the yeast genome. The incongruences in phylogeny and population structure among different genomic regions are likely due, in a large part, to mating between divergent strains. Meiotic products of such hybrids and their subsequent asexual competition can quickly lead to such patterns. Even without meiosis, an apparently rare event in yeast^26^, the production of beneficial aneuploidies during clonal growth can occur, removing some of the chromosomes derived from the hybrid-forming strains.

We identified SNPs associated with variation in growth rate under various environmental stresses compared to a benign condition and validated a subset of them experimentally. Our experiments revealed cases where deleting a gene from a laboratory strain has no appreciable phenotypic effect yet mutations in the gene cause phenotypic differences in natural strains. Because the gene is annotated to be functional in the laboratory strain, our finding suggests that gain-of-function mutations are involved in natural phenotypic variations such that forward genetics can sometimes identify the genetic basis that is invisible by reverse genetics of laboratory strains using gene deletion. Alternatively, our results may indicate the action of epistasis such that a mutation has a phenotypic effect in one but not another genetic background. We also observed that an associated SNP has several underlying causal genetic variants, whose phenotypic effects could even be opposite to one another. These observations add much complexity to association mapping, especially if the goal is to identity causal genetic variants. That we observed all of the above phenomena in the mapping of one trait suggests that this key model system for understanding eukaryotic cell biology still has much to teach us about the genetic basis of phenotypic diversity.

## METHODS

### Strains and strain construction

The strains sequenced in this work were obtained from the authors of two previous studies^12,14^ and are listed in **Supplementary Data 1**. The information of geographic location and environment of each strain is also provided when available. Most strains used are originally diploid and homothallic, and contain no tractable genetic marker, making tracking strains difficult and the maintenance of stable haploid strains, necessary for many studies, impossible. To produce a set of strains useful to the community, we adapted the approach used in the construction of the *S. cerevisiae* gene deletion collection to introduce drug resistance markers flanked by two unique, strain-identifying, 20-nucleotide DNA barcodes at the *HO* (*YDL227C*) locus of each strain (outlined in **Supplementary Fig. 10**). This process simultaneously removed the strains’ ability to mating-type switch and introduced a reliable means for strain tracking. Diploid strains were transformed using the lithium acetate method^39^ with minor alterations. The ~1µg of transforming *HO*-targeting DNA contained a G418 sulfate resistance marker flanked by strain-specific barcodes and was produced by two successive polymerase chain reaction (PCR) amplifications. We first amplified the *KanMX4* cassette from plasmid *pFA6a-KanMX4* ^40^ using two 74-nucleotide primers each containing a unique 20-nucleotide barcode, the sequences necessary for its amplification (U1 + U2 or D1 + D2), and priming sites for the second PCR (**Supplementary Data 8**). The second PCR used a dilution of the product of the first PCR as the template and added sequences homologous to regions upstream and downstream of *HO* for targeting and replacement of the locus (**Supplementary Fig. 10**). The primers used in this PCR differed by strain to maintain lineage-specific SNPs in the region. A full list of the primers used can be found in **Supplementary Data 8**. To ensure that the barcodes assigned to each strain are novel and maintain their compatibility with those in gene deletion strains^4^, MoBY (molecular barcoded yeast) ORF library^41^, and existing technologies used to estimate barcode frequency, we employed unused barcode sequences already present on the widely used Tag4 array^42^. We confirmed successful insertion of the *KanMX4* cassette by PCR and confirmed their sequence using Sanger sequencing.

From these heterozygous *HO* marked diploids (*a/α HO/ho::Uptag-KanMX4-Downtag*), stable haploid strains were obtained by sporulation on potassium acetate media followed by ascus digestion and tetrad dissection. G418 resistant colonies were identified by replication to YPD media containing 300µg/ml G418 sulfate (Gold Biotechnology, US). Colony PCR was used to determine the mating type of individual colonies, and single MAT**a** and MATα colonies were streaked to obtain a pure strain of each mating type. Samples were grown overnight and frozen at -80°C in 20% glycerol for long-term storage. To allow for the easy formation of diploids between any two strains, we switched the drug resistance cassette carried by MAT**a** and MATα strains to hygromycin B and nourseothricin, respectively (Gold Biotechnology). This was achieved by the standard LiAc method using a PCR product produced by the use of primers specific to the TEF promoter and terminator common to all three drug resistance cassettes^40,43^.

Two sets of strains were treated slightly differently due to their genotypes. First, RM11 was previously made into a stable haploid strain by insertion of a *KanMX4* cassette at the *HO* locus, resulting in the deletion of the targeting region we used in all other strains^44^. To insert the appropriate barcodes into this background, unique homologous primers were used to target and replace the existing *KanMX4* cassette with a *HphMX4* marker amplified from plasmid pAG32 ^43^. Unique barcodes were then added and the cassette was switched back to *KanMX4*. Second, three strains (S288c, W303 and RM11) were already heterothallic haploids. After insertion of the barcoded cassette at the *HO* locus, these strains were transformed with plasmid pCM66, which contains a galactose inducible copy of *HO* and a nourseothricin drug resistance marker, to obtain strains of both mating types. After transformation, nourseothricin resistant cells were grown with galactose as the sole carbon source at 30°C without shaking for 8 hours to induce expression of *HO*. This allowed for mating-type switching and subsequent mother-daughter cell mating to produce diploids. Cells were then streaked for single colonies on YPD (1% yeast extract, 2% peptone, 2% glucose, 2% agar) plates, and the ploidy of single colonies was checked by colony PCR using mating-type-specific primers. Diploid colonies were streaked for single colonies on fresh, non-selective, YPD plates and assayed for nourseothricin resistance. A single colony unable to grow in the presence of the drug, and therefore having lost the plasmid, was selected for each strain.

We attempted to produce genetically tractable strains for each of the 85 strains whose genomes we sequenced, but found some to be unamenable to our approach, either due to natural resistance to the drugs used or an inability to successfully sporulate and produce viable offspring of both mating types. The full details of all tractable strains created and the reason for missing strains are outlined in **Supplementary Data 1**.

### Genome sequencing

Each of the 85 strains was streaked from frozen stocks onto YPD plates. Following two days of growth, a single colony was picked into 5ml of liquid YPD media and grown to saturation (36h at 30°C with shaking). Cultures were centrifuged to collect cells, and DNA was extracted using standard methods. Dried DNA pellets were resuspended in 70µl of Tris-EDTA (pH8.0), the DNA was quantified, and the purity was assessed, before DNA storage at -80°C. Illumina libraries were constructed using a protocol modified from a previous study^45^. Briefly, 5µg of genomic DNA was sheared using a Covaris S220 (duty cycle 10%, intensity 4, cycles/burst 200, time 55s), of which 2µg was used in library construction. To select DNA fragments of the desired size range (~400 nucleotides), we used DNA binding Magna beads to perform dual size selection. The fragments were blunt-end repaired, adapter ligated, and nick filled to repair the adapter overhangs. Finally, sequences necessary for multiplexing and cluster formation on an Illumina HiSeq2000 were added by PCR. Equal amounts of each library were combined and run across two paired-end 100-nucletide lanes (43 strains in one lane and 42 in a second) of an Illumina HiSeq2000 at the University of Michigan DNA sequencing core.

### Read mapping and SNP/indel calling

Reads were first trimmed using Cutadapt^46^ to remove adapter sequences. Bowtie2 v2.1.0 ^47^ was used to map reads to the S288c reference (R64-1-1) genome under the sensitive local alignment mode, allowing up to 3 mismatches/indels per read. Pertinent statistics obtained during the mapping process are listed in **Supplementary Data 2**. Paired reads were considered non-concordant and discarded from further analysis if apparent mapping locations were more than 1200 nucleotides apart or if the paired reads appeared to completely overlap one another. Paired reads were also removed from further analysis if either read was found to map ambiguously. Finally, we removed PCR duplicates by discarding all but one copy of any read pair found to map to exactly the same genomic position.

SAMtools v0.1.18 ^48^ and VarScan v2.3.6 ^49^ were used to identify SNPs and indels within each genome. Only variants identified by both programs were used in downstream analysis. To further reduce false calls due to misalignment of reads to the reference genome, we removed variants that showed a significant strand bias (binomial *P* < 0.001) or invariant distance to the end of supporting reads (VDB < 0.0015)^50^. Only the most likely variant is listed for the indel and homozygous SNP lists. For the heterozygous SNP list, maximum likelihood genotype inferred by SAMtools is reported. To reduce errors in estimating allele frequencies, we used only segregating sites with reads covering the variant in each of the 85 strains except when identifying pseudogenizing variants.

### Phylogenetic reconstruction

We reconstructed a maximum composite likelihood neighbor-joining tree using MEGA 5.2 with all homozygous SNPs and all substitution types^51^. We allowed heterogeneous rates amongst lineages and heterogeneous rates amongst sites. Clades were identified in line with previous studies^12,14^. To assess the strength of support for the phylogeny, we performed 1000 bootstraps. Phylogenies of individual chromosomes were reconstructed using the same method.

### Population structure

To assess the population structure of the 190 strains, we used a model-based Markov Chain Monte Carlo (MCMC) algorithm implemented in fastSTRUCTURE^25^. For genome-wide population structure analysis, we randomly selected 10% of non-singleton homozygous SNPs. One hundred runs of fastSTRUCTURE for each of *K* = 2 to 9 were performed, with other parameters set as the default. *K* = 7 was found to be the best. The population structure that exhibited the maximum mean likelihood was plotted using R. Finally, the population structure of each chromosome was determined using all homozygous SNPs on the specific chromosome at *K* = 7.

### Linkage disequilibrium (LD)

LD measured by *r*^2^ between every pair of SNPs was calculated using custom code. We then computed the average *r*^2^ for all SNP pairs with a distance in the range between *x*-99 and *x* nucleotides, where *x* = 100, 200, 300, …, and 200,000. We computed the expected LD between unlinked SNPs by calculating the mean *r*^2^ of 10,000 random pairs of SNPs located on different chromosomes. Following Schachereret et al.^14^, for each distance range, we presented the difference between an observed *r*^2^ and the expected *r*^2^ of unlinked SNPs in Fig. 1d.

### Population genomic analysis of natural selection

We used only SNP sites that are dimorphic and for which the ancestral state could be unambiguously assigned in population genomic analysis. To infer SNP ancestral states, we took advantage of the published orthology information and multi-species genome sequences^52^. We used T-Coffee^53^ and the default settings in BioPerl to align the coding sequences of *S. paradoxus, S. mikatae*, and *S. bayanus* with the orthologous coding sequences of the *S. cerevisiae* reference sequence R64-1-1. Using these multiple-sequence alignments, we considered the states of *S. paradoxus, S. mikatae*, and *S. bayanus* for each SNP site and unambiguously assigned its ancestral state if at least two of these outgroup species were in agreement.

We found polymorphisms that result in nonsense mutations to have much lower derived allele frequencies (DAFs) than nonsynonymous polymorphisms, which in turn have lower DAFs than synonymous polymorphisms (**Supplementary Fig. 3**). This pattern suggests that purifying selection against nonsense mutations is generally stronger than that against nonsynonymous mutations, which is in turn stronger than that against synonymous mutations. To examine whether purifying selection acts on synonymous mutations, especially in genes with strong codon usage bias (CUB), we measured CUB by codon-adaptation index (CAI) of yeast genes previously published^54^. We divided genes into two bins: those with CAI > 0.6 and those with CAI ≤ 0.6. We found that synonymous polymorphisms in high-CAI genes tend to have lower DAFs than those in low-CAI genes (**Supplementary Fig. 3**), supporting the hypothesis of purifying selection against synonymous mutations in genes with strong CUB. For each SNP category, we also calculated the population genetic statistics Tajima’s *D* ^55^, Fu and Li’s *F* ^56^, and Fay and Wu’s *H* ^57^. Compared with synonymous polymorphisms, the more negative values of *D* and *F* for nonsynonymous and nonsense polymorphisms are consistent with the excess of rare alleles, and the less negative values of *H* are consistent with the deficiency of common alleles (**Supplementary Table 1**).

To assess potential positive selection at the protein level, we counted the number of synonymous polymorphisms (*P*_S_), nonsynonymous polymorphisms (*P*_N_), synonymous substitutions between *S. cerevisiae* and *S. paradoxus* (*D*_S_) and nonsynonymous substitutions between *S. cerevisiae* and *S. paradoxus* (*D*_N_) in each gene. The McDonald-Kreitman test^58^ was performed using a two-tailed Fisher’s exact test within R followed by a Bonferroni multiple-test correction. To calculate the proportion of amino acid substitutions driven by positive selection (*α*) for each gene, we used *S. paradoxus* as an outgroup. We first determined if *D*_N_/*P*_N_ > *D*_S_/*P*_S_. When *D*_N_/*P*_N_ > *D*_S_/*P*_S_, we calculated *α* by 1-*D*_S_*P*_N_/(*D*_N_*P*_S_); otherwise, we calculated *α*’ by 1-*D*_N_*P*_S_/(*D*_S_*P*_N_), which represents the fraction of nonsynonymous mutations under purifying selection. We found that the distribution of *α* is largely consistent with widespread purifying selection and relatively few instances of positive selection (**Supplementary Fig. 4**). In addition, McDonald-Kreitman tests of individual genes failed to detect a significant signal of positive selection for any gene after Bonferroni correction. This result is consistent with a previous analysis of a smaller yeast dataset^12^. McDonald-Kreitman tests suggested that 6.9% of genes are under significant purifying selection (Bonferroni corrected *p*-value < 0.05).

Because slightly deleterious alleles can bias the estimation of *α*, we further estimated *α* using the approach proposed by Eyre-Walker and Keightley^59^. Briefly, their approach uses polymorphism data to estimate the distribution of fitness effects of new deleterious mutations (DFE) and then uses DFE to predict the number of neutral and adaptive substitutions. When using this approach with a two-epoch model, we found the estimated *α* to be -0.11 (**Supplementary Table 2**), which is again consistent with the lack of signal of positive selection.

Recently, Messer and Petrov^60^ proposed a heuristic method to estimate *α* by considering α values for polymorphic sites with different levels of DAF. Following Messer and Petrov^60^, we applied the extended version of the McDonald-Kreitman test to all genes. Starting with all SNPs, we sequentially raised the threshold for DAF and recalculated α using only SNPs that pass the threshold. Using MATLAB, we applied a nonlinear least-square method to fit the data to the function of *α*_*x*_ = *a* + *be*^-*cx*^, where *x* is the DAF threshold and α_*x*_ is the corresponding *α*. We restricted *b* < 0 and *c* > 0. While the theoretical support of Messer and Petrov’s approach is lacking, this approach, surprisingly, converges on an estimated value of 0.55 for *α*, suggesting that ~55% of between-species sequence divergence at nonsynonymous sites is due to positive selection (**Supplementary Fig. 5a**). Interstingly, the plot of *α* vs DAF showed a clear valley around intermediate DAFs that has not been observed previously. Suspecting the reason might be heterogeneous selection across strains, we partitioned strains based on phylogenetic clustering. We found that if we seperated strains from the Wine/European cluster from all other strains and performed the same analysis on the two groups separately, the plots of *α* vs DAF were dramatically different from each other. The extrapolated α values were -0.36 for the Wine/European cluster and 0.56 for all other strains. This difference in the estimate of *α* was not caused by sampling in general, but the specific partitions chosen (**Supplementary Fig. 5b, c**). In addition, the *α* vs. DAF plot for strains not in the Wine/European cluster showed a better fit to an exponential curve than the combined analysis (adjusted *R*^2^ = 0.85 without Wine/Europen strains vs. adjusted *R*^2^ = 0.69 for all strains). Overall, these results suggest that the historical action of natural selection within these two groups has been different and that positive selection in yeast may be more common than initially expected, especially outside of domesticated wine strains. These results, however, are inconsistent with the results obtained from using the method of Eyre-Walker and Keightley^59^, where no positive values are apparent when using either only strains in the Wine/European cluster or only strains outside the Wine/European cluster (**Supplementary Table 2**).

### De novo assembly and identification of non-reference genes

*De novo* genome assembly was performed using SOAPdenovo2 v2.04 ^61^ with *K* = 51 for all adapter-trimmed reads from each of the 85 strains we sequenced. Basic statistics of the genome assemblies are listed in **Supplementary Data 3**. To examine the quality of the assemblies, we used BLASTn to search for the *KanMX4* vector sequence present at the *HO* locus of 81 genomes. We then used it as an anchor to extract the strain-specific barcodes (UPTAG and DSTAG). We successfully recovered the strain-specific barcodes for all 81 genomes. To rule out false positive *de novo* gene calls, Exonerate v2.2.0 ^62^ was used to align known genes in the reference genome of S288c to the assembled contigs. Known genes were localized onto contigs by prioritizing the Exonerate hits by (i) best hit with syntenic neighbor genes at either side, (ii) best hit with 100% query sequence coverage, and (iii) hits longer than 200 nucleotides that do not overlap with better hits by more than 30 nucleotides - in case a gene is split among multiple contigs. Having identified the locations of known genes on *de novo* contigs, we used GeneMarkS v4.17 ^63^ to perform gene predictions and compared these to the locations of known genes.

Predicted genes that showed no overlap with known gene locations were considered candidate non-reference genes. To avoid false positives caused by un-localized known genes, we used BLASTn to align the predicted genes with cDNA sequences of known genes. All predicted genes with hits covering 80% of the query, or with a > 200-nucleotide region that is > 90% identical with any reference gene, were removed. In order to classify the origin of remaining non-reference genes, we retrieved the best hit in the NCBI “nr” database reported by BLASTn and tBLASTx for each non-reference gene. If the best hit was in the reference genome, it was also removed unless there was a premature stop codon for the hit region in the reference genome. Finally, we removed non-reference genes with best hits to sequences derived from vectors, synthetic constructs, phages, bacterial genomes, or tandem elements (**Supplementary Data 4**).

To access the expression levels of the non-reference genes, RNA-seq data of 23 *S. cerevisiae* strains generated by SOLiD were downloaded^29^. The color space RNA-seq reads from each strain were mapped to known and predicted non-reference genes using bowtie^64^, allowing up to 2 mismatches in color space. The best hits for each read were then used to calculate the Reads Per Kilobase of transcript per Million mapped reads (RPKM) for each gene.

### Phenotypic consequences of deleting non-reference genes

Two newly identified non-reference genes were deleted from the genetic backgrounds in which they were discovered using the G418 sulfate resistance cassette *KanMX4* that was PCR-amplified from plasmid *pFA6a-KanMX4* ^40^, followed by PCR confirmation (see **Supplementary Data 8** for primer sequences). *Non-Ref-1* was deleted from strain UWOPS87-2421, while *Non-Ref-2* was deleted from strain CLIB272. Both backgrounds were MATa *ho::NatMX4* genotypes detailed below.

To test the phenotypic consequences of the deletion of these two genes, growth curves were obtained using a Bioscreen C (Growth Curves, USA). *Non-Ref-1* deletion strain and wild-type strain were grown in Complete Supplemented Media (CSM) containing varying concentrations (0, 3, 6, 12.5, 25, 37.5, 50, 75, and 100µg/ml) of fluconazole (Sigma Aldrich). *Non-Ref-2* deletion strain and wild-type strain were also grown in CSM media but using varying concentrations (0, 1.5, 3, 6, 12, 25, 50, and 100µM) of catharidin. To initiate the growth, we grew 5ml CSM cultures from frozen stocks for 24h. Each strain was diluted 200 times into 350µl of the appropriate drug-containing media in triplicate. Growth curves were collected at 30°C for 60h using the Wide-Band 420-580nm filter every 20 minutes. Maximum growth rate (OD/h) was collected from growth curve data as in a previous study^11^ and efficiency (Max OD) was calculated by taking the average of the 3^rd^-7^th^ highest OD values recorded. Two-tailed *t-*tests were used to assess the significance of differences between genotypes. Three replications were performed for each growth curve.

### Identification of intron losses

Based on S288c reference genome annotation, we built a sequence database containing all known exon-exon junctions up to 101 nucleotides from each side of the junction. To search for reads supporting an intron loss event, all reads were mapped to this database by Bowtie2. We required at least 95% coverage of the query read and that the read mapped to at least 20 nucleotides in each of two adjacent exons. We further filtered ambiguous mappings on the reference genome by BLASTn with e-value cutoff at 0.01. To search for reads supporting the presence of introns, the same procedure was conducted for all sequences annotated as exonintron borders in the S288c reference genome. Finally, intron loss was declared in strains in which at least two reads span the exon-exon junction but no read spans the two corresponding exon-intron borders.

### Identifying potential copy number variants (CNVs) and aneuploidies

To assess gene duplication/deletion events, read pairs that were concordantly mapped to the reference genome following filtering of potential PCR duplicates were analyzed using Cufflinks v2.1.1 ^65^ to generate Fragments Per Kilobase of transcript per Million mapped reads (FPKM) values for each coding sequence (CDS). Potential CNVs of individual genes, as well as aneuploidies and large scale duplications, were then identified by dividing each FPKM by that obtained from the same CDS in the reference strain S288c.

### Simultaneous phenotyping of barcoded yeast strains

To phenotype all strains with unique barcodes, we used bar-seq^23^ to estimate their relative growth rates in each of seven environments. The barcoded strains were mixed approximately equally and combined with the diploid homozygous gene deletion collection (Invitrogen). Each non-deletion strain was present at approximately twice the initial population size of each gene deletion strain. The initial pool of strains was grown for approximately two generations in 25ml of YPD media at 30°C before the resulting culture, termed generation 0, was used to initiate competitions in each of the experiments. To reduce the effect of genetic drift, large populations were maintained throughout competitions with regular transfers to fresh media every 4-5 generations to maintain populations in exponential growth. Populations were competed for approximately 30 generations (6 transfers) and samples were stored at -80°C following each transfer. Following preliminary investigations, we chose to carry out in-depth analysis of the populations following the second transfer (~10 generations) in YPD at 30°C, YPD at 40°C, YPD + 1.25M NaCl at 30°C, YPD + 8% EtOH at 30°C, YPD + 4mM paraquat (superoxide) at 30°C, YPD + 3mM Hydrogen peroxide at 30°C, and YPD + 1mM cobalt chloride at 30°C, respectively.

To determine the frequency of strains in the pooled population at a given time point, we extracted genomic DNA from samples using a Puregene Yeast/Bacteria DNA extraction kit (Qiagen). DNA barcodes were amplified by PCR using Accuprime pfx (Invitrogen). The primers used for barcode amplification also added sequences necessary for cluster formation and sequencing primer annealing on the Illumina platform. Because the downstream barcode is known to be missing in some deletion strains^66^, only the upstream barcodes were used. Fifty base-pair single-end sequence reads were obtained using one lane of an Illumina Genome Analyzer IIx at the University of Michigan DNA Sequencing Core. The Illumina Pipeline software version 1.6 was used for base calling from the image data. Because all sequences started with the same 18 base pairs of the PCR primer region and this uniformity adversely affected base calling, we removed the first 18 sequencing cycles before base calling. We used the previously published “gene-barcode map”^67^ with the addition of our own strains’ barcode identities to assign each read to a particular strain allowing for only a single mismatch. We followed a previously published outline of bar-seq analysis^68^. We required that barcodes be represented by at least 40 total counts in populations analyzed before (generation 0) and after competition. Count numbers were then normalized using the TMM method implemented in the R package egdeR^69^. Because we are interested in phenotypes that are specific to a given condition, as opposed to the general fitness across environments, we computed fitness in a specific condition relative to that in YPD (benign condition). Overall there were 79 natural *S. cerevisiae* strains and 4498 deletion strains with usable data.

Let the number of reads for a genotype of interest at the beginning of the competitions be *N*_0_ and the corresponding number for the reference genotype (i.e., the strain lacking *HO*) be *M*_0_. Let the numbers of reads for the above two genotypes after competition in the benign environment of YPD at 30°C be *N*_1_ and *M*_1_, respectively, and the corresponding numbers in a stressful environment of interest be *N*_2_ and *M*_2_, respectively. The competition lasted for *g* = ~10 generations in each environment. Note that the inaccuracy in *g* does not affect comparison of relative fitness among strains, because the same *g* applies to all strains. The fitness of the genotype of interest relative to that of the reference genotype in the stressful environment, relative to that in the benign environment is 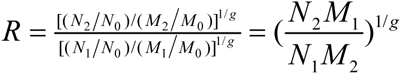.

Two biological replicates of the competition and bar-seq were conducted in YPD at 30°C. Based on the two estimates of *N*_1_/*M*_1_, we tested whether *N*_2_/*M*_2_ is significantly different from *N*_1_/*M*_1_ (i.e., whether *R* is significantly different from 1), followed by FDR correction for multiple testing.

## GWAS

We followed a multistep GWAS approach^70,71^ outlined in **Supplementary Fig. 12**. For each environment, we first scaled and centered the relative growth rates. We removed all sites with a minor allele frequency below 5% across the set of strains for which we had phenotype data. We then removed sites for which data were missing in >5% of strains and all mitochondrial sites. The remaining 123,121 sites were converted into a 0, 0.5, 1 format to represent homozygous non-reference, heterozygous, and homozygous reference states in each strain, respectively.

Besides true positives, individual SNPs can be significantly associated with a phenotype because of chance or the confounding effect of population structure. Because over- or under-correction of population structure can lead to a loss of statistical power or false positives, respectively, we performed the GWAS in a number of steps to attempt to find an appropriate balance. First, we performed an association between the normalized phenotype values and each SNP without controlling for population structure by a simple linear regression method without covariates. Second, we used this unstructured association to rank all SNPs on the basis of their statistical significance of association. From this list, we performed a series of associations by maximum likelihood using EMMA^72^. To control for population structure, we estimated a kinship matrix based on a specific set of SNPs. To define this set, we started with the 1000 SNPs most significantly associated with the phenotype in the unstructured analysis and then successively added the next 1000 most significant SNPs from the unstructured analysis until 120,000 SNPs were included. For each association, the genomic control factor, lambda, was calculated using gcontrol2 within R ^73^. We identified the minimal kinship set that controlled for population structure based on where lambda first hit 1, or if it failed to do so, was minimized (**Supplementary Fig. 12**). Third, we ran an additional series of associations centered on the 3000 SNP region identified by lambda using kinship sets in 50 SNP windows. Again, we found the smallest kinship set where lambda hit 1, or was at its minimum. Finally, we performed an association for the 500 most significant SNPs from the unstructured association. In each case, we used the estimated kinship set that was optimal for lambda, minus any SNPs within 10 kb of the focal SNP, to estimate population structure. Any variant in this final association with a *p-*value below 0.0001 (i.e., 5%/500) was classified as significant. For these SNPs, we identified the coding region nearest to the variant as well as its immediate neighbors. These candidate SNPs generally cover a 4-10kb region which is approximately 4-8 times the range over which LD is seen to break down.

### Experimental validation of GWAS results

To confirm the validity of our GWAS approach and to narrow down the genes responsible for the observed phenotypic variations within the regions surrounding the significant SNPs, we performed reciprocal hemizygosity tests^24^. We chose to concentrate on a subset of SNPs significantly associated with relative fitness at 40°C. The same candidate genes were deleted from haploid *MATa ho::HygMX4* backgrounds that showed a high relative fitness and a low relative fitness using a *KanMX4* marker. Appropriate strain hybrids were formed by mating of these deletion strains to *MATα* cells. Two types of hybrid were formed to allow for the relative fitness of reciprocal hemizygotes to be ascertained. The first was formed by mating of deletion-carrying haploids to *MATα ho::NatMX4* of the other genetic background and the second was formed by mating to *MATα ho::TDH3p-YFP-NatMX4* cells. This resulted in four hybrid strains for each gene to be tested, fluorescent and non-fluorescent hybrids lacking the high-fitness allele (ΔH-F and ΔH-NF) and fluorescent and non-fluorescent hybrids lacking the low-fitness allele (ΔL-F and ΔL-NF). All strains were frozen at -80°C until needed. To perform competitive fitness assays, freezer samples were inoculated into 1ml of YPD and grown with shaking for 24 hours at 30°C. Diploid hybrids were paired so that each competition included two hybrids carrying reciprocal deletions of the alternate alleles and identifying markers. That is, ΔH-F was paired with ΔL-NF in the first competition, and ΔL-F was paired with ΔH-NF in the second competition. Competitions were performed by mixing equal volumes of the two strains to form the *t*_0_ population which was diluted 1000 fold into 1.5ml of fresh YPD media pre-warmed at 40°C. Strains were competed for 36 hours, or ~10 generations, at 40°C with shaking till saturation (*t*_1_). All competitions were performed in 96 deep-well plates with 4 replicates of each competition. The frequencies of fluorescent and non-fluorescent cells within *t*_0_ and *t*_1_ populations were determined by flow cytometry using a BD Accuri C6 attached to a hypercyt sampling robot. After data cleaning to remove artifacts, the relative fitness of the competing strains was calculated using 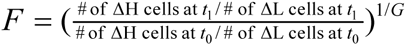, where *G* = 10 denotes the number of generations of competition. The same experiment was repeated in YPD at 30°C. Relative fitness was computed by dividing fitness at 40°C by fitness at 30°C. Five replicate competitions were performed for each set of strains and the relative fitness was calculated by averaging over the reciprocal YFP-marked strain pairs for each gene. Statistical significance was obtained by *t* tests.

### Data access

The yeast genome sequences determined in this work have been deposited to NCBI with the BioProject ID of PRJNA320792.

## Acknowledgements

We thank Gianni Liti and Leonid Kruglyak for sharing the yeast strains that are sequenced in this study and Audrey Gasch for help accessing read data from previously sequenced strains. This work was supported by a research grant from the U.S. National Science Foundation (MCB-1329578) to J.Z.

## References

1. Goffeau, A. et al. Life with 6000 genes. Science 274, 546, 563–7 (1996).

2. Hillenmeyer, M. E. et al. The chemical genomic portrait of yeast: uncovering a phenotype for all genes. Science 320, 362–5 (2008).

3. Ohya, Y. et al. High-dimensional and large-scale phenotyping of yeast mutants. Proc. Natl. Acad. Sci. U. S. A. 102, 19015–20 (2005).

4. Giaever, G. et al. Functional profiling of the Saccharomyces cerevisiae genome. Nature 418, 387–91 (2002).

5. Douglas, A. C. et al. Functional analysis with a barcoder yeast gene overexpression system. G3 2, 1279–89 (2012).

6. Sopko, R. et al. Mapping Pathways and Phenotypes by Systematic Gene Overexpression. Mol. Cell 21, 319–330 (2006).

7. Kellis, M., Patterson, N., Endrizzi, M., Birren, B. & Lander, E. S. Sequencing and comparison of yeast species to identify genes and regulatory elements. Nature 423, 241–254 (2003).

8. Hittinger, C. T. Saccharomyces diversity and evolution: a budding model genus. Trends Genet. 29, 309–17 (2013).

9. Scannell, D. R. et al. The Awesome Power of Yeast Evolutionary Genetics: New Genome Sequences and Strain Resources for the Saccharomyces sensu stricto Genus. G3 1, 11–25 (2011).

10. Dujon, B. et al. Genome evolution in yeasts. Nature 35–44 (2004).

11. Warringer, J. et al. Trait variation in yeast is defined by population history. PLoS Genet. 7, e1002111 (2011).

12. Liti, G. et al. Population genomics of domestic and wild yeasts. Nature 458, 337–41 (2009).

13. Wang, Q.-M., Liu, W.-Q., Liti, G., Wang, S.-A. & Bai, F.-Y. Surprisingly diverged populations of Saccharomyces cerevisiae in natural environments remote from human activity. Mol. Ecol. 21, 5404–17 (2012).

14. Schacherer, J., Shapiro, J. a, Ruderfer, D. M. & Kruglyak, L. Comprehensive polymorphism survey elucidates population structure of Saccharomyces cerevisiae. Nature 458, 342–5 (2009).

15. Cromie, G. A. et al. Genomic Sequence Diversity and Population Structure of Saccharomyces cerevisiae Assessed by RAD-seq. G3. 3, 2163–71 (2013).

16. Hose, J. et al. Dosage compensation can buffer copy-number variation in wild yeast. Elife 4, e05462 (2015).

17. Strope, P. K. et al. The 100-genomes strains, an S. cerevisiae resource that illuminates its natural phenotypic and genotypic variation and emergence as an opportunistic pathogen. Genome Res. (2015).

18. Bergström, A. et al. A High-Definition View of Functional Genetic Variation from Natural Yeast Genomes. Mol. Biol. Evol. msu037- (2014).

19. Gonçalves, M. et al. Distinct Domestication Trajectories in Top-Fermenting Beer Yeasts and Wine Yeasts. Curr. Biol. 26, 2750–2761 (2016).

20. Gallone, B. et al. Domestication and Divergence of Saccharomyces cerevisiae Beer Yeasts. Cell 166, 1397–1410.e16 (2016).

21. Connelly, C. F. & Akey, J. M. On the prospects of whole-genome association mapping in Saccharomyces cerevisiae. Genetics 191, 1345–53 (2012).

22. Diao, L. & Chen, K. C. Local ancestry corrects for population structure in Saccharomyces cerevisiae genome-wide association studies. Genetics 192, 1503–11 (2012).

23. Smith, A. M. et al. Quantitative phenotyping via deep barcode sequencing. Genome Res. 19, 1836–1842 (2009).

24. Steinmetz, L. M. et al. Dissecting the architecture of a quantitative trait locus in yeast. Nature 416, 326–30 (2002).

25. Raj, A., Stephens, M. & Pritchard, J. K. fastSTRUCTURE: variational inference of population structure in large SNP data sets. Genetics 197, 573–89 (2014).

26. Tsai, I. J., Bensasson, D., Burt, A. & Koufopanou, V. Population genomics of the wild yeast Saccharomyces paradoxus: Quantifying the life cycle. Proc. Natl. Acad. Sci. U. S. A. 105, 4957–62 (2008).

27. Juneau, K., Miranda, M., Hillenmeyer, M. E., Nislow, C. & Davis, R. W. Introns regulate RNA and protein abundance in yeast. Genetics 174, 511–8 (2006).

28. Parenteau, J. et al. Deletion of many yeast introns reveals a minority of genes that require splicing for function. Mol. Biol. Cell 19, 1932–41 (2008).

29. Skelly, D. A. et al. Integrative phenomics reveals insight into the structure of phenotypic diversity in budding yeast. Genome Res. 23, 1496–504 (2013).

30. Gbelska, Y., Krijger, J.-J. & Breunig, K. D. Evolution of gene families: the multidrug resistance transporter genes in five related yeast species. FEMS Yeast Res. 6, (2006).

31. Menichetti, F. et al. High-dose fluconazole therapy for cryptococcal meningitis in patients with AIDS. Clin. Infect. Dis. 22, 838–40 (1996).

32. Martin, M. V. The use of fluconazole and itraconazole in the treatment of Candida albicans infections: a review. J. Antimicrob. Chemother. 44, 429–37 (1999).

33. Jungwirth, H., Wendler, F., Platzer, B., Bergler, H. & Högenauer, G. Diazaborine resistance in yeast involves the efflux pumps Ycf1p and Flr1p and is enhanced by a gain-of-function allele of gene YAP1. Eur. J. Biochem. 267, 4809–16 (2000).

34. Brôco, N., Tenreiro, S., Viegas, C. A. & Sá-Correia, I. FLR1 gene (ORF YBR008c) is required for benomyl and methotrexate resistance in Saccharomyces cerevisiae and its benomyl-induced expression is dependent on pdr3 transcriptional regulator. Yeast 15, 1595–608 (1999).

35. Lissina, E. et al. A systems biology approach reveals the role of a novel methyltransferase in response to chemical stress and lipid homeostasis. PLoS Genet. 7, e1002332 (2011).

36. Bloom, J. S., Ehrenreich, I. M., Loo, W. T., Lite, T.-L. V. & Kruglyak, L. Finding the sources of missing heritability in a yeast cross. Nature 494, 234–7 (2013).

37. Winzeler, E. a. Functional Characterization of the S. cerevisiae Genome by Gene Deletion and Parallel Analysis. Science 285, 901–906 (1999).

38. He, X., Qian, W., Wang, Z., Li, Y. & Zhang, J. Prevalent positive epistasis in Escherichia coli and Saccharomyces cerevisiae metabolic networks. Nat. Genet. 42, 272–6 (2010).

39. Cubillos, F. a, Louis, E. J. & Liti, G. Generation of a large set of genetically tractable haploid and diploid Saccharomyces strains. FEMS Yeast Res. 9, 1217–25 (2009).

40. Wach, A., Brachat, A., Pöhlmann, R. & Philippsen, P. New heterologous modules for classical or PCR-based gene disruptions in Saccharomyces cerevisiae. Yeast 10, 1793–808 (1994).

41. Ho, C. H. et al. A molecular barcoded yeast ORF library enables mode-of-action analysis of bioactive compounds. Nat. Biotechnol. 27, 369–77 (2009).

42. Pierce, S. E. et al. A unique and universal molecular barcode array. Nat. Methods 3, 601–3 (2006).

43. Goldstein, A. L. & McCusker, J. H. Three new dominant drug resistance cassettes for gene disruption in Saccharomyces cerevisiae. Yeast 15, 1541–53 (1999).

44. Brem, R. B., Yvert, G., Clinton, R. & Kruglyak, L. Genetic dissection of transcriptional regulation in budding yeast. Science (80-.). 296, 752–5 (2002).

45. Rohland, N. & Reich, D. Cost-effective, high-throughput DNA sequencing libraries for multiplexed target capture. Genome Res. 22, 939–46 (2012).

46. Martin, M. Cutadapt removes adapter sequences from high-throughput sequencing reads. EMBnet.journal 17, 10–12 (2011).

47. Langmead, B. & Salzberg, S. L. Fast gapped-read alignment with Bowtie 2. Nat. Methods 9, 357–9 (2012).

48. Li, H. et al. The Sequence Alignment/Map format and SAMtools. Bioinformatics 25, 2078–2079 (2009).

49. Koboldt, D. C. et al. VarScan 2: Somatic mutation and copy number alteration discovery in cancer by exome sequencing. Genome Res. 22, 568–76 (2012).

50. Daneck, P. et al. High levels of RNA-editing site conservation amongst 15 laboratory mouse strains. Genome Biology 13, R26 (2012).

51. Tamura, K. et al. MEGA5: Molecular evolutionary genetics analysis using maximum likelihood, evolutionary distance, and maximum parsimony methods. Mol. Bio. Evol. 28, 2731–9 (2011).

52. Scannell, D. R. et al. The Awesome Power of Yeast Evolutionary Genetics: New Genome Sequences and Strain Resources for the Saccharomyces sensu stricto Genus. G3. 1, 11–25 (2011).

53. Notredame, C., Higgins, D. G. & Heringa, J. T-Coffee: A novel method for fast and accurate multiple sequence alignment. J. Mol. Biol. 302, 205–217 (2000).

54. Qian, W., Yang, J.-R., Pearson, N. M., Maclean, C. & Zhang, J. Balanced codon usage optimizes eukaryotic translational efficiency. PLoS Genet. 8, e1002603 (2012).

55. Tajima, F. Statistical method for testing the neutral mutation hypothesis by DNA polymorphism. Genetics 123, 585–595 (1989).

56. Fu, Y. X. & Li, W. H. Statistical tests of neutrality of mutations. Genetics 133, 693–709 (1993).

57. Fay, J. C. & Wu, C. I. Hitchhiking under positive Darwinian selection. Genetics 155, 1405–1413 (2000).

58. McDonald, J. H. & Kreitman, M. Adaptive protein evolution at the Adh locus in Drosophila. Nature 351, 652–4 (1991).

59. Eyre-Walker, A. & Keightley, P. D. Estimating the rate of adaptive molecular evolution in the presence of slightly deleterious mutations and population size change. Mol. Biol. Evol. 26, 2097–108 (2009).

60. Messer, P. W. & Petrov, D. A. Frequent adaptation and the McDonald-Kreitman test. Proc. Natl. Acad. Sci. U. S. A. 110, 8615–20 (2013).

61. Luo, R. et al. SOAPdenovo2: an empirically improved memory-efficient short-read de novo assembler. Gigascience 1, 18 (2012).

62. Slater, G. S. C. & Birney, E. Automated generation of heuristics for biological sequence comparison. BMC Bioinformatics 6, 31 (2005).

63. Besemer, J., Lomsadze, A. & Borodovsky, M. GeneMarkS: a self-training method for prediction of gene starts in microbial genomes. Implications for finding sequence motifs in regulatory regions. Nucleic Acids Res. 29, 2607–2618 (2001).

64. Langmead, B., Trapnell, C., Pop, M. & Salzberg, S. L. Ultrafast and memory-efficient alignment of short DNA sequences to the human genome. Genome Biol 10, R25 (2009).

65. Trapnell, C. et al. Transcript assembly and quantification by RNA-Seq reveals unannotated transcripts and isoform switching during cell differentiation. Nat. Biotechnol. 28, 511–5 (2010).

66. Deutschbauer, A. M. et al. Mechanisms of haploinsufficiency revealed by genome-wide profiling in yeast. Genetics 169, 1915–25 (2005).

67. Qian, W., Ma, D., Xiao, C., Wang, Z. & Zhang, J. The genomic landscape and evolutionary resolution of antagonistic pleiotropy in yeast. Cell Rep. 2, 1399–410 (2012).

68. Robinson, D. G., Chen, W., Storey, J. D. & Gresham, D. Design and Analysis of Bar-seq Experiments. G3 (Bethesda). 4, 11–8 (2014).

69. Robinson, M. D., McCarthy, D. J. & Smyth, G. K. edgeR: a Bioconductor package for differential expression analysis of digital gene expression data. Bioinformatics 26, 139–40 (2010).

70. Listgarten, J. et al. Improved linear mixed models for genome-wide association studies. Nature Methods 9, 525–526 (2012).

71. Listgarten, J., Lippert, C. & Heckerman, D. FaST-LMM-Select for addressing confounding from spatial structure and rare variants. Nat. Genet. 45, 470–1 (2013).

72. Kang, H. M. et al. Efficient control of population structure in model organism association mapping. Genetics 178, 1709–1723 (2008).

73. Devlin, B. & Roeder, K. Genomic control for association studies. Biometrics 55, 997–1004 (1999).

